# Stress during puberty and adulthood pregnancy impact histone acetylation regulators in the hypothalamus

**DOI:** 10.1101/2024.12.01.626244

**Authors:** Laiklyn A.M. Luther, Samantha L. Higley, Kathleen E. Morrison

## Abstract

Undergoing stressful events during puberty puts women at risk for a variety of negative outcomes, and this risk is heightened if they become pregnant later in life. We previously demonstrated that stress during puberty combined with pregnancy in adulthood led to a blunted response of the hypothalamic-pituitary-adrenal stress axis in humans and mice. We have begun to understand the mechanisms underlying this effect by examining the paraventricular nucleus of the hypothalamus (PVN), a key regulator of the HPA axis. Prior studies uncovered an increase in chromatin openness within the PVN of the at-risk mice, with bioinformatic analyses implicating histone acetylation in this increased openness. Here, we measured the activity of histone acetyltransferase (HATs) and histone deacetylase (HDACs), the writers and erasers of histone acetylation, within the PVN to further characterize how stress during puberty and pregnancy may be interacting to produce a blunted stress response. We found that histone acetylation tone within the PVN is predictive of prior transcriptional and chromatin results, with pregnant, pubertally stressed females having a pro-acetylation tone within the PVN. These findings establish a role for regulators of acetylation in the open chromatin landscape characteristic in the PVN of pregnant, pubertally stressed females. Overall, this study provides insight into the epigenetic mechanisms underlying female-relevant risk for stress dysregulation, a central endophenotype of affective disorders.

## Introduction

Undergoing adverse childhood experiences (ACEs) heightens the risk for a variety of negative outcomes in adulthood (Rosenberg et al., 2007; Merrick et al., 2017). For women, exposure to stress during adolescence results in vulnerability to developing affective disorders later in life. We previously demonstrated that undergoing stress during the critical developmental window of puberty resulted in a blunted stress response amongst female mice and humans, but only during pregnancy (Morrison et al., 2017, 2020a; Duffy et al., 2024), which is a period of dynamic hormonal and neuroanatomical change (Servin-Barthet et al., 2023). This is consistent with other work showing an association between ACE exposure and negative outcomes during the perinatal period, including psychosocial difficulty (Racine et al., 2018), suicidal ideation (Zhong et al., 2016), parental distress and anxiety (Zak-Hunter et al., 2023), and prenatal substance use (Hemady et al., 2022). Clinicians have advocated for expanding our understanding of how the life history of a woman impacts her mental and physical health both during the perinatal period and throughout the lifespan. Although we have evidence that multiple life experiences, such as pubertal stress and pregnancy, compound across the lifespan to influence risk for negative outcomes, we understand less about the biological mechanisms underpinning this association.

We have previously shown that stress during puberty has a lasting effect on the physiology and behavior of mice and humans. Peripartum mice and humans with a history of pubertal stress exposure show a blunted hormonal stress response, which was associated with increased postnatal depression scores in humans (Morrison et al., 2017). Our subsequent work focused on the paraventricular nucleus of the hypothalamus (PVN), the central regulator of the hypothalamic-pituitary-adrenal (HPA) axis-driven hormonal stress response. We established that pubertal stress has lasting effects on the epigenome of the PVN of mice. Specifically, ATAC-sequencing data showed that there were significantly more open regions in the chromatin within PVN of female mice that undergo stress during puberty, but only when they are pregnant in adulthood (Morrison et al., 2020a). This pregnancy-state dependent change in the chromatin landscape of pubertally-stressed females mirrors our prior findings in the PVN transcriptome and stress axis responding. This may be due to the fact that chromatin state has a critical impact on transcriptional activity (D’Oliveira Albanus et al., 2021). There are several mechanisms by which the chromatin landscape can be altered to either promote or repress gene expression. One way is through post-translational modification (PTM) of the tails of histones, which are the proteins around which DNA is wrapped. There are several types of PTMs, in which specific amino acids in the tails of histone proteins undergo modifications (Millán-Zambrano et al., 2022). The addition of acetyl groups, primarily to the amino acid lysine, is a key mechanism that leads to increased openness of chromatin and enhanced gene expression. Histone acetyltransferases (HATs) and histone deacetylases (HDACs) are classes of enzymes that regulate acetylation. HATs ‘write’ acetyl marks by catalyzing the addition of acetyl marks to lysine residues, while HDACs ‘erase’ acetyl marks (Yang and Seto, 2007). Both enzymes work to control acetylation and ultimately represent potent regulators of the chromatin landscape.

Whether our chromatin findings are due to the bioinformatically-predicted changes in acetylation are unknown. Individually, early life stressors and pregnancy have been associated with changes to histone acetylation in various brain regions. Early life stressors such as maternal separation produce lasting effects on histone acetylation within the ventral tegmental area, forebrain, hippocampus, and amygdala (Levine et al., 2012; Xie et al., 2013; Blaze et al., 2015; Seo et al., 2016; Xu et al., 2022; Geiger et al., 2024). Regulation of acetylation in brain regions like the medial preoptic area and amygdala of female mice are important in supporting normative maternal behavior (Stolzenberg et al., 2014; Mayer et al., 2019; Seward et al., 2022). However, it is unknown if undergoing stress during puberty and pregnancy during adulthood together result in any synergistic effect on histone acetylation and whether the hypothalamus is subject to the same kind of reprogramming as these previously examined brain regions.

Here, we measured the activity of regulators of acetylation in the PVN at several time points in the lifespan: before, immediately after, and long after pubertal stress. By using this approach, we aimed to get a sense of overall acetylation potential of the chromatin within the PVN. Specifically, we investigated HAT and HDAC activity in the PVN of female and male mice during both adolescence and adulthood. In adult females, we included pregnancy as a factor, as we have seen the highest risk for HPA axis dysfunction when pubertally stressed females become pregnant later in life. Ultimately, our findings give insight into the molecular mechanisms involved in the open chromatin landscape of pubertally-stressed, pregnant female mice. Understanding this process is critical to understanding why females have an increased risk for certain disorders across the lifespan.

## Experimental Procedures

### Animals

All mice used were of an in-house mixed C57BL/6:129 strain. Any mouse used for breeding was virgin at the time of breeding. Food and water were available to animals *ad libitum* and animals were kept on a 12-hour light/dark cycle (lights on at 8:00am). All procedures were approved by the West Virginia University Institutional Animal Care and Use Committee and were conducted in accordance with the National Institutes of Health Guide for the Care and Use of Laboratory Animals.

### Pubertal stress

Beginning at postnatal day (PN) 21 (**Figure 1**), mice were singly housed and underwent 14 days of chronic variable stress (CVS) (Morrison et al., 2016, 2017, 2020a; Gautier et al., 2024). Two stressors were administered per day beginning within the first hour of lights on (on at 8:00 am) during the daily nadir of circadian corticosterone levels. CVS lasted two hours each day. Two stressors were assigned to be applied simultaneously each day and varied between three different sensory modalities: olfactory (puma odor [1:200 2-Phenethylamine, CAS 64-04-0 in mineral oil]; 70% ethanol), auditory (white noise or owl screech at 50-80 db), and tactile (novel object exposure [8 toy glass marbles in cage], wire mesh, wet bedding, no bedding, multiple cage changes [3 within 2 hours], and 15 minute acute restraint). Mice experienced rotating combinations of stressors (tactile and auditory, tactile and olfactory, or auditory and olfactory), combinations of which also rotated between specific stressors each day. After the last stress exposure on PN34, mice were pair-housed with a same-sex, same-treatment littermate. Control mice were weaned from dams at PN28 and housed with a same-sex cage mate. Mice in the CVS group were weaned at PN21 to include early neglect as a component of the stressor experience, consistent with our previous work in this model (Morrison et al., 2016, 2017, 2020a; Gautier et al., 2024).

**Figure 1.**
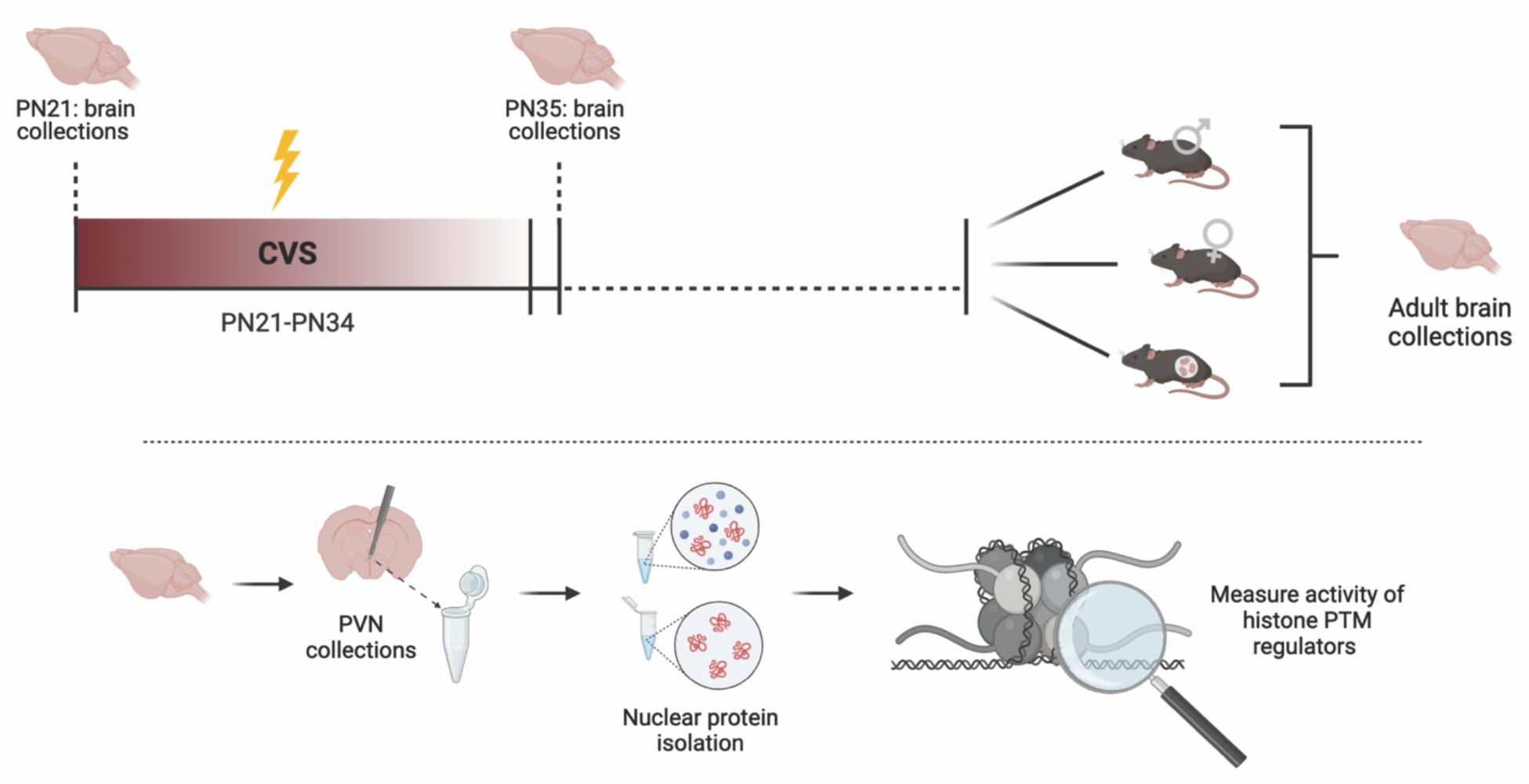
Timeline of experimental procedures. Animals underwent CVS from PN21-34. During this, two stressors which target different sensory modalities were administered for two hours (except for restraint stressor which lasted 15 minutes) each morning. Brains were collected prior to stress (PN21) and 24 hours after last stressor (PN35). In adulthood, brains were collected from control and CVS males, females, and pregnant females during late pregnancy (17.5 dpc). PVN tissue was isolated and prepared for histone PTM regulator measurement. CVS = chronic variable stress, PN = postnatal day, dpc = days post conception, PVN = paraventricular nucleus of the hypothalamus, PTM = post-translational modification. Created with BioRender.com.

### Breeding scheme

At 9-11 weeks of age, female mice were bred with naïve male mice. Breeders were paired together for 1-3 nights. Pregnancy was confirmed through copulation plug check, and pregnant females were moved to single housing and left undisturbed until euthanization at 17.5 days post conception (dpc).

### Tissue collection

Brains were collected in the morning (lights on) at one of several time points: PN21 (prior to stress), PN35 (24 hours after the final stressor), and adulthood (11-14 weeks). At PN21, groups included control male and female mice. PN35 groups included control and CVS males and females. Adulthood groups included control and CVS males, females, and late pregnant females. Late pregnant females were collected on the morning of 17.5 dpc. For collection, mice were anesthetized with 5% isoflurane prior to decapitation. Brain tissue was dissected and stored immediately on dry ice prior to being stored at -80°C until later use. To obtain a tissue sample of the PVN, brains were cryosectioned at -20°C. Once the correct anatomical region was identified using Paxinos and Franklin stereotaxic coordinates, two 300 um slices were taken. The PVN was biopsied from those slices using a pre-chilled 1.0 mm hollow biopsy tool. Biopsies were placed in a pre-chilled 1.5 mL tube and stored at -80°C. To ensure that all biopsies captured the PVN, 10 um slices taken immediately before and after slicing the region of interest were stained using neutral red dye. Slices were imaged and analyzed to confirm that each sample included the PVN. If a sample did not meet the anatomical criteria, it was not included in analysis.

### Protein isolation

Protein was isolated from the PVN samples using the Epiquik Nuclear Extraction Kit, per manufacturer instructions (Epigentek, OP-0002). Protein concentrations were measured using the Qubit Protein Assay Kit (ThermoFisher, Q33211).

### Histone enzyme activity quantification

HAT activity was measured using the EpiQuik HAT Activity/Inhibition Assay Kit (Epigentek, P-4003). HDAC activity was measured using the Epigenase HDAC Activity/Inhibition Direct Assay Kit (Epigentek, P-4034). Assays were conducted and optical density (OD) values for all assays were obtained via a microplate reader per manufacturer instructions. OD values were used to calculate enzyme activity for each sample (OD/hour/mg protein), and relative activity was calculated for statistical analysis (relative activity = activity for sample / average activity for age-matched Female Controls). HAT/HDAC balance was calculated by dividing HAT activity by HDAC activity for a given sample, and relative balance was calculated for statistical analysis (relative balance = balance for sample / balance for age-matched female controls).

### Statistical analyses

The relative activity of each enzyme type (HAT, HDAC), as well as the balance of activity (HAT/HDAC) was analyzed using the most appropriate approach for the question. Data were considered outliers if they were ± two standard deviations from the mean and were excluded from analysis. Unpaired t-tests were used to analyze sex differences at PN21. Outcomes at PN35 were analyzed via two-way analysis of variance (ANOVA) with sex and pubertal stress (CVS) as factors. Adult outcomes were analyzed via two-way ANOVA with sex/pregnancy and pubertal stress (CVS) as factors. Tukey post-hoc tests were utilized following any significant interaction. All statistical analyses were conducted using Prism (Graphpad) with alpha for significance set at 0.05. We considered alpha levels < 0.1 but > 0.05 to be trending effects. Group sizes for each outcome (**Table S1**) and output of all individual statistical tests can be found in the **Supplemental Material**.

## Results

### Pubertal stress impacts acetylation tone in the adult PVN

We have previously shown that pubertal stress (CVS) resulted in an altered chromatin landscape in the PVN that was characterized by increased openness of chromatin (Morrison et al., 2020a). An open chromatin landscape in the PVN has been associated with negative outcomes, including a dysfunctional stress response. Bioinformatic analyses indicated that histone acetylation was the posttranslational modification that could be responsible for this open chromatin and permissive gene expression. Here, we tested whether CVS had a lasting effect on the regulators of histone acetylation in the PVN by measuring baseline HAT and HDAC activity and assessing the balance of activity of these classes of enzymes.

For HAT activity (**Figure 2A**), there was a significant CVS x sex/pregnancy status interaction (F(2, 57) = 3.30 *p* = 0.04, **Table S2**). Tukey post-hoc tests showed that none of the individual groups were significantly different from each other (*p* > 0.05), although examination of the data indicates that the interaction stems from a finding that CVS males had a pattern of reduced HAT activity. Indeed, males exposed to CVS had 19% less HAT activity in the PVN than control males, while CVS induced a 7% reduction in non-pregnant females and a 5% increase in pregnant females. This effect is reflected by the trend for a main effect of CVS (F(1, 57) = 3.007, *p* = 0.09).

**Figure 2.**
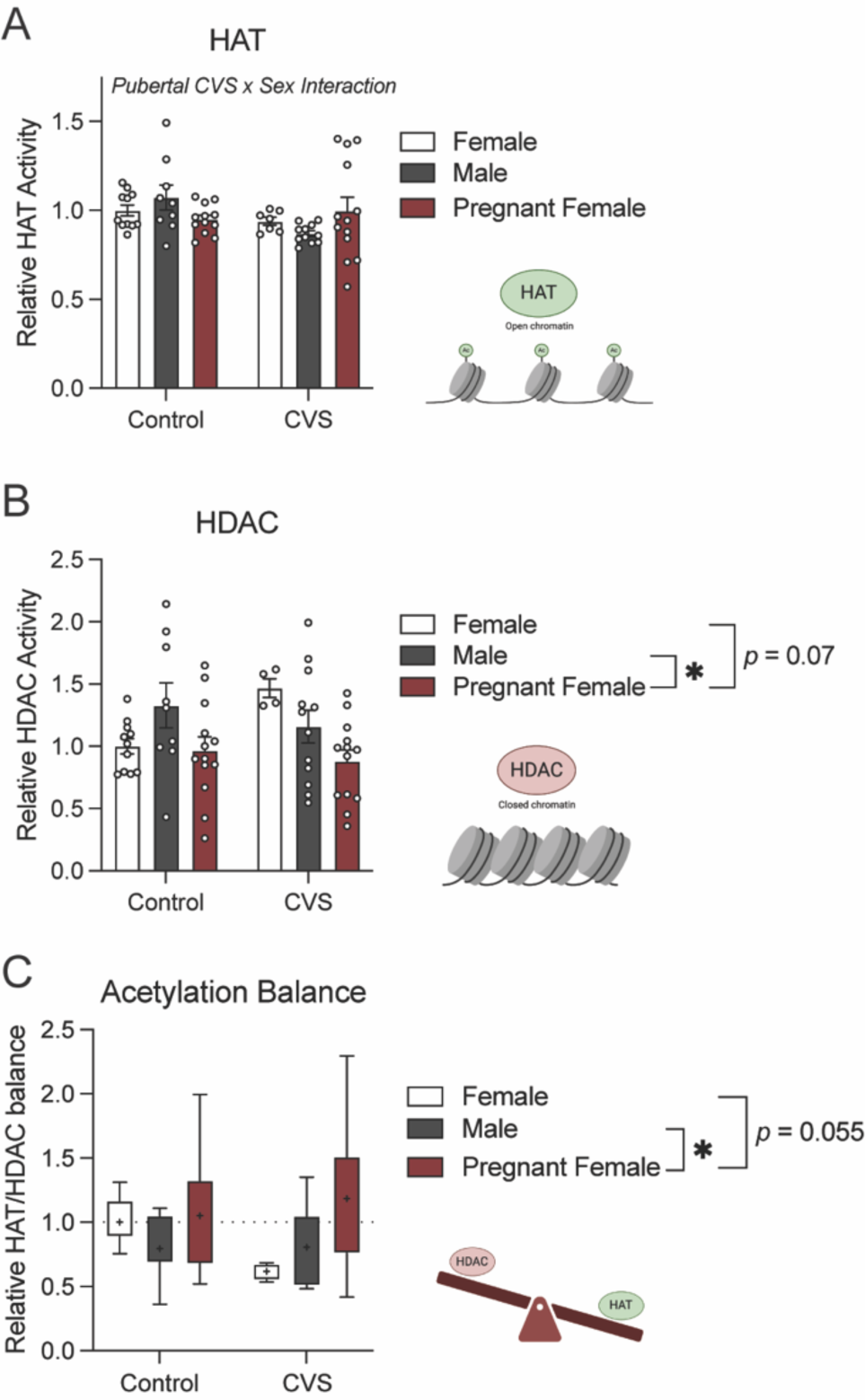
Pubertal stress and sex/pregnancy status interact to influence activity of histone acetylation regulators in the PVN of adults. (**A**) HAT enzymes are associated with open chromatin and increased transcription. Pubertal stress and sex/pregnancy interacted to influence HAT activity in the PVN. Post-hoc analysis showed no statistical differences between groups, although CVS adult males tend to have decreased HAT activity compared to control adult males. (**B**) HDAC enzymes are associated with closed chromatin and repression of transcription. There was a main effect of sex/pregnancy on HDAC activity in the PVN. There was significant difference between male and pregnant females, such that pregnant females had decreased HDAC activity. There was a non-significant trend for a difference between non-pregnant females and pregnant females. (**C**) The HAT/HDAC balance indicates the overall acetylation tone that would result from the opposing actions of HATs and HDACs. Values > 1 indicate more openness, while values < 1 indicate a tone that supports closed chromatin. There was a main effect of sex/pregnancy on HAT/HDAC balance in the PVN. Pregnant females had significantly higher HAT/HDAC balance compared to males, with a trend for a difference between control females and pregnant females. The finding that pregnant, CVS adult females have a PVN acetylation tone that would lead to open chromatin is consistent with our previous sequencing and transcriptomic findings. HAT = histone acetyltransferase, HDAC = histone deacetylase, PVN = paraventricular nucleus of the hypothalamus, \**p* < 0.05 on Tukey’s HSD post-hoc test. + indicate mean value on box plot graphs. All values are relative to within-measure Female Controls. Data are mean ± SEM.

There was a significant main effect of sex/pregnancy status on HDAC activity (**Figure 2B**, F(2, 56) = 4.83, *p* = 0.01). Regardless of CVS exposure, pregnant females had significantly lower HDAC activity in the PVN than did adult males (*p* = 0.02) and a trending difference from adult non-pregnant females (*p* = 0.07). There was no difference between the two groups that are ‘resilient’ to stress axis dysfunction, adult non-pregnant females and males (*p* > 0.99). There was a trend for an interaction between CVS and sex/pregnancy status (F(2, 56) = 2.79, *p* = 0.07), which is likely driven by the impact of pregnancy on HDAC activity in the PVN of adult, CVS females. Pregnancy had a minimal impact on control females, who display 4% less HDAC activity in the PVN when pregnant compared to non-pregnant controls. In contrast, pregnancy had a more prominent impact on CVS females, who displayed a 59% decrease in HDAC activity in the PVN when pregnant.

Finally, we examined the overall histone acetylation tone to predict whether the PVN was in an ‘open’ vs ‘closed’ chromatin state. We calculated the ratio of HAT to HDAC activity within a sample (**Figure 2C**). There was a main effect of sex/pregnancy status (F(2, 52) = 4.93, *p* = 0.01). Post-hoc testing revealed that pregnant females had overall increased HAT/HDAC balance in the PVN compared to males (*p* = 0.02) and a trend towards increased HAT/HDAC balance compared to non-pregnant females (*p* = 0.055). There was no difference in HAT/HDAC balance between males and non-pregnant females (*p* > 0.99). Overall, the group with the highest risk for dysfunction of the stress axis – pregnant, CVS females – had the highest HAT/HDAC tone (18% higher than control Females and 58% higher than CVS, nonpregnant females). The finding that the average expression is towards more activity of acetyl writers, which are associated with openness, is consistent with our prior ATAC-Seq findings of the highest chromatin openness in this group.

### Pubertal stress does not impact acetylation tone in the adolescent PVN

We measured baseline activity of acetylation regulators before and after the pubertal CVS exposure. There were no sex differences in baseline HAT activity (**Figure 2A**), HDAC activity (**Figure 2C**), or HAT/HDAC balance (**Figure 2E**) in the PVN at PN21 (all *p* > 0.05, **Table S3**). Therefore, when males and females entered CVS on PN21, there were no differences in the activity of these enzymes or their relative balance of activity in the PVN that might indicate a preexisting risk or resilience factor. We also examined baseline activity of these regulators twenty-four hours following the end of CVS, at PN35. There were no significant effects of sex or CVS on baseline HAT activity (**Figure 2B**), HDAC activity (**Figure 2D**), or HAT/HDAC balance (**Figure 2F**) at PN35 (all *p* > 0.05, **Table S4**). Although not significant, there was some variability in HAT/HDAC balance by sex, with males showing a 5% reduction (more ‘closed’ tone) and females showing a 17% increase (more ‘open’ tone) in response to pubertal stress.

*Pubertal stress alters the developmental trajectory of acetylation regulators in the PVN* We compared the relative HAT, HDAC, and HAT/HDAC balance across all ages and states (**Figure 4**). We set all within-age measures to be relative to age-matched control females, which allowed us to examine levels of acetylation associated with risk versus resilience for pubertal stress-induced dysfunction. Pubertally-stressed females are at the most risk for lasting negative outcomes during stress, and they display the highest pro-acetylation HAT/HDAC balance at two key time points: twenty-four hours after the end of CVS and during late pregnancy. In contrast, males are the most resilient to the impacts of pubertal stress. This is mirrored in their HAT/HDAC balance, in which males display the lowest HAT/HDAC balance, indicative of an anti-acetylation tone.

**Figure 3.**
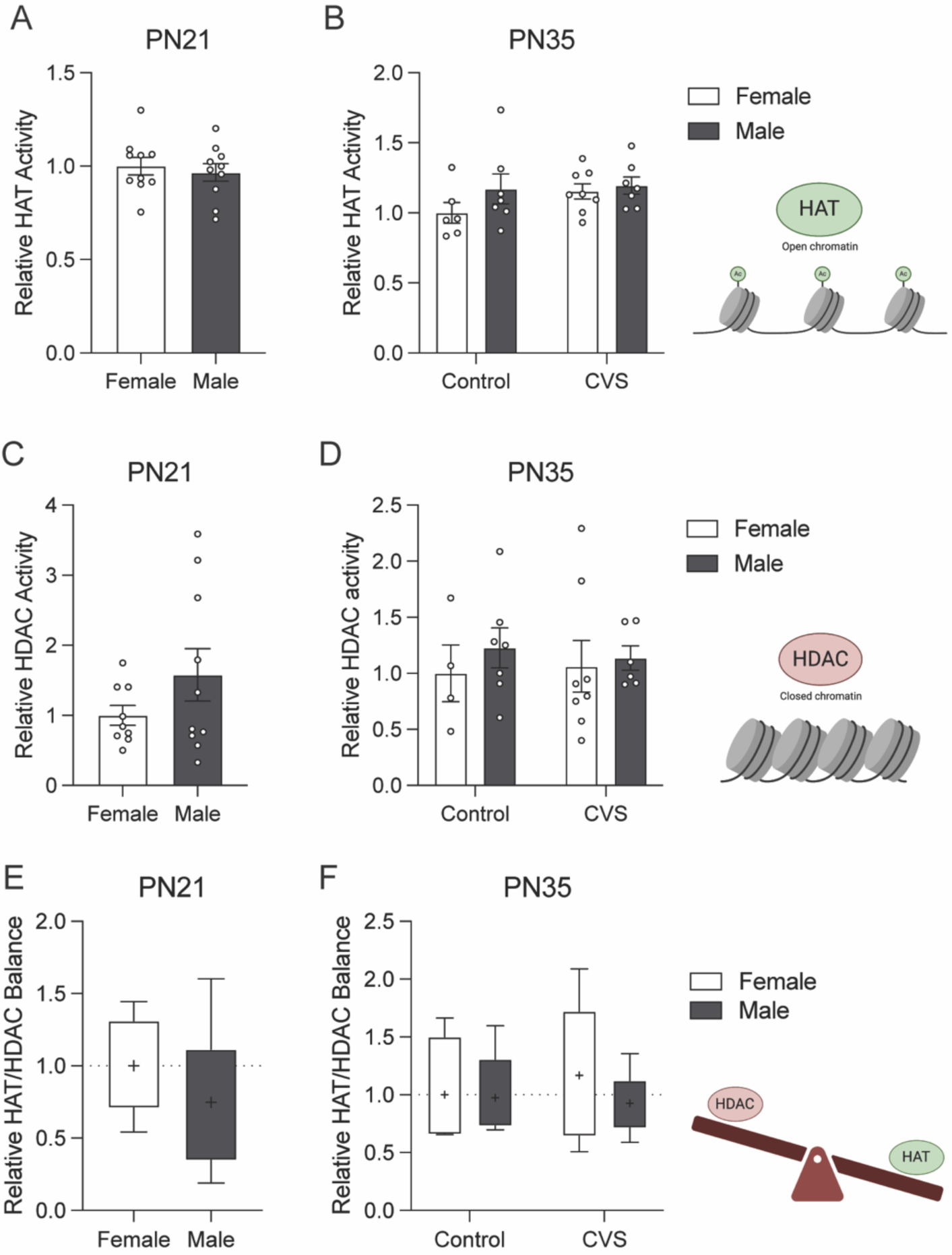
There was no impact of pubertal stress or sex on the activity of histone acetylation regulators in the PVN of adolescents. (**A,C,E**) The activity of HAT and HDAC enzymes, and the HAT/HDAC balance were measured at PN21, prior to any stressor exposure. (**B,D,F**) These same outcomes were measured at baseline at PN35, 24 hours after the end of pubertal stress. **(A)** HAT enzymes are associated with open chromatin and increased transcription. Prior to stress, there was no difference in HAT activity in the PVN between males and females. **(B)** At PN35, there was no effect of pubertal stress or sex on HAT activity in the PVN. **(C)** HDAC enzymes are associated with closed chromatin and repression of transcription. There was no difference in HDAC activity between males and females in the PVN prior to beginning stress. **(D)** At PN35, there was no effect of pubertal stress or sex on HDAC activity in the PVN. **(E)** The HAT/HDAC balance indicates the overall acetylation tone that would result from the opposing actions of HATs and HDACs. Values > 1 indicate more openness, while values < 1 indicate a tone that supports closed chromatin. When looking at HAT/HDAC balance in the PVN, there was no difference between males and females at PN21. **(F)** There was also no effect of pubertal stress or sex on HAT/HDAC balance at PN35. HAT = histone acetyltransferase, HDAC = histone deacetylase, PVN = paraventricular nucleus of the hypothalamus. + indicate mean value on box plot graphs. All values are relative to within-age, within-measure Female Controls. Data are mean ± SEM.

**Figure 4.**
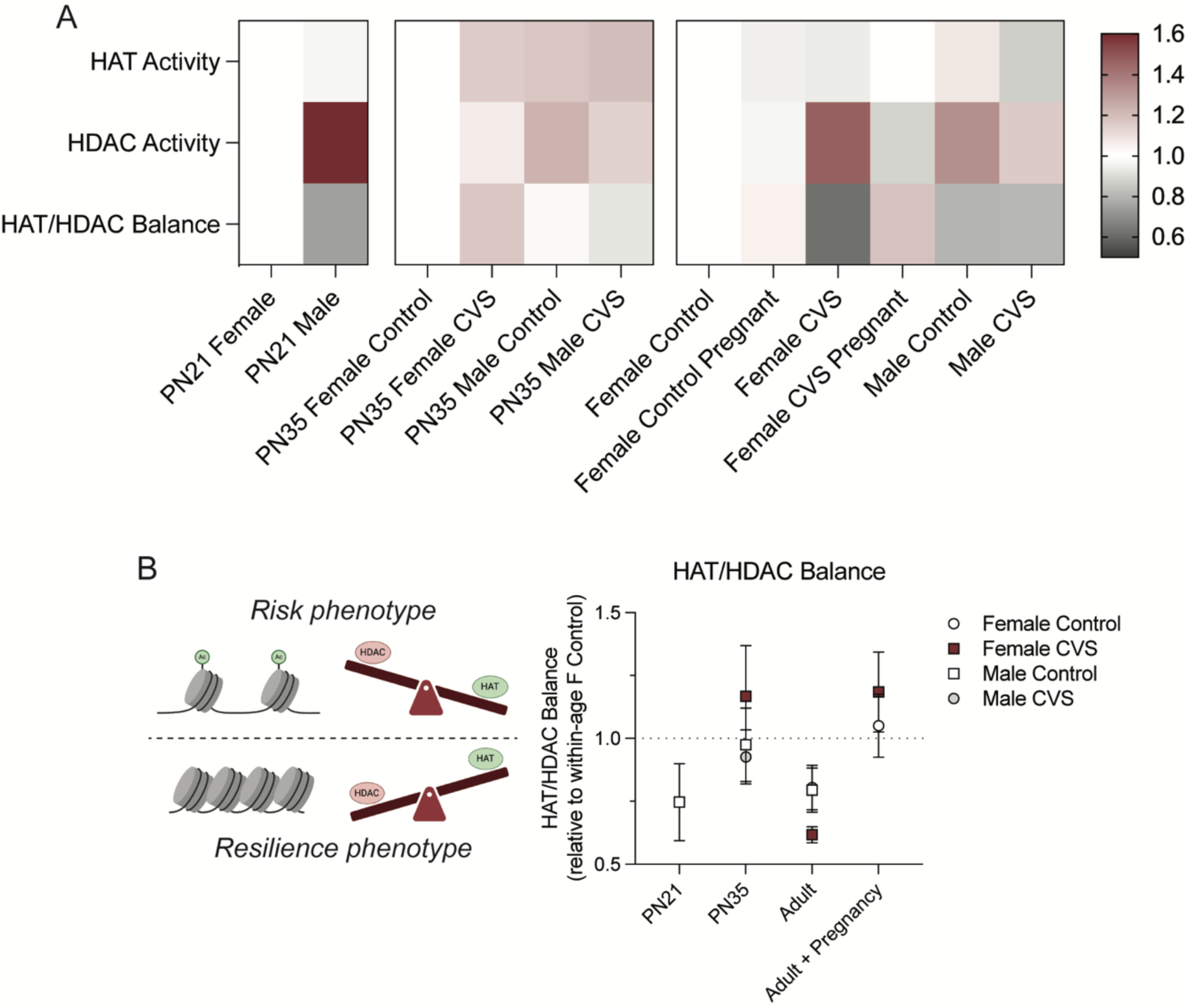
Changes in acetylation tone align with prior findings of open chromatin in the PVN of pubertally-stressed, pregnant females. (**A**) Heatmaps showing the relative activity to age-matched female control for HAT and HDAC enzymes and the relative HAT/HDAC balance at each time point (PN21, PN35, adulthood) for each group. Red indicates increased activity or balance, and gray indicates decreased activity or balance. (**B**) In adulthood, pregnant CVS females show a HAT/HDAC balance that is indicative open chromatin. A more open chromatin landscape is associated with a blunted stress response (risk phenotype), while more closed chromatin may indicate resilience. Animals that are unlikely to show an altered stress response – control females, nonpregnant CVS females, and males – all show a HDAC predominated tone, which is associated with closed chromatin. Schematic elements created with BioRender.com. PVN = paraventricular nucleus of the hypothalamus, PN = postnatal day, HAT = histone acetyltransferase, HDAC = histone deacetylase. All values are relative to within-age, within-measure Female Controls.

## Discussion

We previously established that stress during puberty combined with pregnancy in adulthood led to a blunted HPA axis response in female mice and humans, a phenotype associated with affective dysregulation (Naughton et al., 2014; Morrison et al., 2017).

Further investigation showed that increased chromatin openness within the PVN may contribute to this response, and bioinformatic analysis of ATAC-seq data implicated histone acetylation in the increased openness (Morrison et al., 2020a). Acetylation is a post-translational modification that is characterized by the addition of an acetyl group to the tail of a histone and is associated with increased transcriptional activity (Yang and Seto, 2007; Millán-Zambrano et al., 2022). HATs and HDACs are enzyme classes that play an important role in this process by attaching and removing acetyl marks from the tails of histones, respectively. To gain insight into the possible mechanisms underlying the phenotype we previously observed, we sought to characterize the activity of these histone PTM regulators in the PVN of mice and whether activity is altered by pubertal stress, sex, and pregnancy in adulthood. Using our translationally relevant mouse model of pubertal adversity, we measured the activity of HATs and HDACs within the PVN at varying time points throughout the mouse lifespan, including during pregnancy in adulthood. In doing so, we found that females who experienced pubertal stress and pregnancy as adults had the most pro-acetylation tone within the PVN, which promotes a more open chromatin landscape.

In our model, we consistently find that exposure to pubertal stress and then pregnancy in adulthood leads to reliable physiological, transcriptional, and epigenetic reprogramming. The differences between pubertally stressed and control females are not observed until they become pregnant in adulthood. It is the exposure to the dynamic experience of pregnancy that is key to uncover the latent physiological, transcriptional, and epigenetic reprogramming. Here, we investigated whether this phenomenon of cumulative stress, where a ‘second hit’ challenge uncovers latent reprogramming (Kuhn et al., 2016), extends to the regulators of acetylation. Pregnant females who underwent stress during puberty displayed a reduction in HDAC activity within the PVN compared to nonpregnant, pubertally-stressed females, suggesting that pubertal stress and pregnancy may interact to produce an inhibition of HDAC activity in the PVN. A decrease in activity of HDACs, which erase acetyl marks, could lead to more acetylation and more open chromatin. We also analyzed the balance between the activity of the opposing mechanisms of HATs and HDACs to understand the overall acetylation tone likely produced in the PVN. We found that pregnant females had a pro-acetylation balance, such that there was a higher HAT tone within the PVN of these animals. This further supports our hypothesis that the activity of HATs and HDACs may contribute to a more open chromatin landscape. These findings suggest that the pro-acetylation tone in pregnant, pubertally stressed females may be driven by the disruption of HDAC activity. Alterations of HAT and HDAC activity in the hippocampus and amygdala have been associated with both risk and resilience for behavior dysfunction following a single prolonged stressor in adolescence (Xu et al., 2022). Future work, including pharmacological manipulation of HATs and HDACs, will allow us to fully determine the specific contributions of these enzymes in the mechanisms of pubertal stress-induced physiological dysfunction.

To understand the developmental origins of the adult phenotypes, we examined HAT and HDAC activity both prior to stress exposure and twenty-four hours following end of stress. Although we did not observe any significant differences in activity or balance, there was some variability during adolescence that may merit further investigation. Notably, pubertally stressed females have an increased, albeit nonsignificant, HAT/HDAC balance, indicating a pro-acetylation tone in the PVN. This effect has been returned to a latent state by adulthood, where HDAC tone predominates in non-pregnant females. Future studies could address whether the days following pubertal stress represent a window of plasticity where either pharmacological or environmental factors could prevent the lasting effects of pubertal stress. This pattern of transient disruption following stress that has become latent in adulthood matches our findings on the impact of pubertal stress on expression of key genes within the PVN (Gautier et al., 2024). These key genes include six immediate early genes, which are permissively expressed in baseline, non-stimulated conditions in the PVN of pregnant, pubertally stressed females. While we have not directly connected the transcriptome and epigenome here, prior work supports such a hypothesis. For instance, maternal separation stress has been shown to increase both histone acetylation and immediate early gene expression within the hippocampus of mice (Xie et al., 2013). Future work will further strengthen this potential mechanistic link between the chromatin landscape, downstream gene expression, and ultimately a dysfunctional stress response.

The question pursued here, and the mouse model used to pursue it, are inherently focused on times in the lifespan when females are vulnerable to the impacts of adversity. Nonetheless, we continue to include male mice in our experiments as an important control, to provide further insight into known sex differences in responses to stressors throughout the lifespan, and to understand one potential aspect of resilience (Rincón-Cortés et al., 2019; Morrison et al., 2020b). In our recent examination of the impact of pubertal stress on the transcriptome of the PVN, we found that baseline mRNA expression in males was less sensitive in adulthood than in females (Gautier et al., 2024). Here, we find that there were significant differences between males and pregnant females in HAT/HDAC balance, with males having a strong HDAC-biased anti-acetylation tone in the PVN. This increased HDAC activity and overall anti-acetylation tone in the PVN may be a protective factor in males. This is consistent with prior work showing that maternal separation stress leads to a decrease in forebrain HDAC activity, increased acetylation, and increased anxiety-like behavior of male mice, and that pharmacological increasing HDAC activity reversed these effects (Levine et al., 2012). In another study, early life maltreatment led to altered histone acetylation of specific genes in the medial prefrontal cortex of female, but not male, mice (Blaze et al., 2015). Altogether, these findings supporting a potential mechanistic role for HDAC enzymes in the lasting sex-specific effects of early life stress.

Our investigation of acetylation was focused on overall activity of HATs and HDACs, which represent two broad classes of acetylation regulators. This approach allowed us to gain a holistic view of the acetylation state within the PVN. However, it will be beneficial to follow up on these findings and focus on specific histone modifications associated with more open chromatin. For example, H3K27ac is an acetyl mark know to be indicative of active enhancers and increased transcription (Ye et al., 2020). Previous work has indicated increases in transcriptional activity, as six immediate early genes were upregulated in the PVN of pubertally-stressed, pregnant female mice (Morrison et al., 2020a). More closely examining specific epigenetic marks will provide a more detailed understanding of the exact mechanisms at play contributing to blunted stress response and ultimately an increased risk for disorder across the lifespan.

Overall, we demonstrate that females who undergo stress during puberty and then become pregnant as adults show the most pro-acetylation tone within the PVN, which indicates increased risk for HPA axis dysfunction. Results of this study provide us with novel insight into how acetylation may be playing a role in the chromatin permissiveness, which we hypothesize is contributing to the altered HPA axis response. Understanding the underlying mechanisms leading to chromatin landscape changes is essential to understanding why females are at a greater risk for certain neuropsychiatric disorders. Knowing what is driving this increased risk will allow for future development of intervention and treatment of negative outcomes.

## Author Contributions

KEM designed experiments. LAML and SLH conducted experiments. LAML and KEM sorted and analyzed data. LAML wrote an initial draft of the manuscript, which was edited to a final version in collaboration with KEM.

## Acknowledgments

This work was supported by National Institutes of Health Grant R00 HD091376 (KEM) and start-up funds to KEM. LAML was additionally funded by the West Virginia University Research Apprenticeship Program and West Virginia University Summer Undergraduate Research Experience program.

## Data Availability

Data will be publicly available via the Open Science Framework repository upon publication.

