## Supplemental Material for "Stress during puberty and adulthood pregnancy impact histone acetylation regulators in the hypothalamus"

### Supplemental Tables

**Table S1.** Final sample sizes by group

| Group | Sample size | Number of litters |
|---|---|---|
| PN21 | | |
| Female | 9-10 | 5-6 |
| Male | 9-10 | 5 |
| PN35 | | |
| Female Control | 4-6 | 3-5 |
| Female CVS | 8 | 5 |
| Male Control | 6-7 | 5-6 |
| Male CVS | 6-7 | 4 |
| Adult | | |
| Female Control Not Pregnant | 11 | 7 |
| Female Control Pregnant | 12-13 | 7-8 |
| Female CVS Not Pregnant | 4-7 | 4-6 |
| Female CVS Pregnant | 12-13 | 6 |
| Male Control | 8-9 | 7-8 |
| Male CVS | 11-12 | 5 |

*Sample size represents HAT, HDAC, and HAT/HDAC measures. CVS = chronic variable stress, PN = postnatal day*

**Table S2.** Full statistical reporting from 2-way ANOVA analysis for adult PVN acetylation regulators

| Measure | Factor | F value | P value | Significance |
|---|---|---|---|---|
| HAT Activity | | | | |
| | Pubertal CVS | F (1, 57) = 3.007 | P=0.088 | ns |
| | Sex/Pregnancy | F (2, 57) = 0.003 | P=0.997 | ns |
| | **Interaction** | F (2, 57) = 3.295 | P=0.044 | * |
| HDAC Activity | | | | |
| | Pubertal CVS | F (1, 56) = 0.432 | P=0.513 | ns |
| | **Sex/Pregnancy** | F (2, 56) = 4.829 | P=0.012 | * |
| | Interaction | F (2, 56) = 2.792 | P=0.0698 | ns |
| HAT/HDAC Balance | | | | |
| | Pubertal CVS | F (1, 52) = 0.591 | P=0.446 | ns |
| | **Sex/Pregnancy** | F (2, 52) = 4.929 | P=0.011 | * |
| | Interaction | F (2, 52) = 1.989 | P=0.147 | ns |

*Measure: activity or balance relative to Female Control Not Pregnant; ns = not significant, $p < 0.05$; Pubertal CVS (Control, CVS); Sex/Pregnancy (Female, Male, Pregnant Female)*

**Table S3.** Results of t-testing for analysis of sex effect on PVN acetylation regulators at PN21

| Measure | t ratio | df | P value | Significance |
|---|---|---|---|---|
| HAT Activity | 0.536 | 18 | 0.599 | ns |
| HDAC Activity | 1.383 | 17 | 0.184 | ns |
| HAT/HDAC Balance | 1.351 | 16 | 0.195 | ns |

*Measure: activity or balance relative to Female;*
*ns = not significant*

**Table S4.** Full statistical reporting from 2-way ANOVA analysis for PN35 PVN acetylation regulators

| Measure | Factor | F value | P value | Significance |
|---|---|---|---|---|
| HAT Activity | | | | |
| | Pubertal CVS | F (1, 24) = 1.369 | P=0.253 | ns |
| | Sex | F (1, 24) = 1.958 | P=0.175 | ns |
| | Interaction | F (1, 24) = 0.719 | P=0.405 | ns |
| HDAC Activity | | | | |
| | Pubertal CVS | F (1, 21) = 0.005 | P=0.946 | ns |
| | Sex | F (1, 21) = 0.513 | P=0.482 | ns |
| | Interaction | F (1, 21) = 0.132 | P=0.720 | ns |
| HAT/HDAC Balance | | | | |
| | Pubertal CVS | F (1, 20) = 0.100 | P=0.755 | ns |
| | Sex | F (1, 20) = 0.507 | P=0.485 | ns |
| | Interaction | F (1, 20) = 0.333 | P=0.570 | ns |

*Measure: activity or balance relative to Female Control; Pubertal CVS (Control, CVS); Sex (Female, Male); ns = not significant*
